# Proton pump inhibitors modulate esophageal epithelial barrier function and crosstalk with eosinophils

**DOI:** 10.1101/2024.08.22.609219

**Authors:** Ravi Gautam, Megha Lal, Margaret C. Carroll, Zoe Mrozek, Tina Trachsel, Jarad Beers, Melanie A. Ruffner

**Affiliations:** Division of Allergy and Immunology, Children’s Hospital of Philadelphia; Department of Pediatrics, Perelman School of Medicine at University of Pennsylvania; Division of Allergy, University Children’s Hospital Zurich, Zurich, Switzerland

**Keywords:** eosinophilic esophagitis, proton pump inhibitor, epithelial barrier, eosinophil, chemokine

## Abstract

**Background:** Eosinophilic esophagitis (EoE) is a chronic allergic disease characterized by esophageal dysfunction, type-2 inflammation, and esophageal eosinophilic infiltrate. While proton pump inhibitor (PPI) therapy is commonly used for EoE management, the underlying mechanism of action remains unclear.

**Methods:** Air-liquid interface culture of esophageal epithelial cells was employed to investigate the impact of the PPI omeprazole on barrier integrity in IL-13-treated cultures. Epithelial chemokine secretion was assessed following stimulation with IL-13 and omeprazole, and the migration of eosinophils from healthy human donors was evaluated using 3 µm pore-sized transwells. A co-culture system of epithelial cells and eosinophils was employed to study chemokine secretion and eosinophil adhesion and activation markers.

**Results:** Omeprazole treatment in the IL-13-treated air-liquid interface (ALI) model resulted in 186 differentially expressed genes and restored barrier integrity compared to ALI treated with IL-13 alone. Omeprazole treatment reduced STAT6 phosphorylation, downregulated calpain 14, and upregulated desmoglein-1 in the IL-13-treated air-liquid interface samples. IL-13-induced upregulation of Eotaxin-3, CXCL10, and periostin, but this was downregulated by omeprazole. Further, the expression of CD11b, CD18, and CD69 was lower on eosinophils from omeprazole-treated epithelial-eosinophil co-cultures, which also had lower levels of eotaxin-3, CXCL10, CCL2, and CCL4.

**Conclusion:** Omeprazole reduced the effects of IL-13 in both the epithelial air-liquid interface model and eosinophil-epithelial co-cultures, reducing barrier dysfunction, chemokine expression, and upregulation of eosinophil adhesion markers.

## INTRODUCTION

Eosinophilic esophagitis (EoE) is a chronic immune-mediated disorder triggered by specific food antigens and is characterized by eosinophil-rich multicellular inflammation and epithelial changes (1). Treatment options for EoE include dietary therapy, pharmacological therapy, or a combination of diet and medication, including proton pump inhibitors (PPIs), corticosteroids, and biologic therapy(2).

PPIs have historically been used as acid suppressants for a variety of disorders like reflux and peptic ulcer disease. PPIs are delivered as a prodrug that becomes activated by gastric acid, inhibits parietal gastric H^+^, K^+^-ATPase, and reduces gastric acid secretion (3). Previously, PPI treatment was considered critical for distinguishing GERD from EoE, but a trial of PPI therapy is no longer deemed necessary to diagnose EoE (4). This change is due to studies showing that PPI treatment resolved the type 2 inflammatory signature in esophageal biopsy tissue similarly to topical corticosteroids(5).

Studies have increasingly focused on how PPIs reduce eosinophilia and inflammation in EoE (6–8). Several pathways have been implicated, including inhibition of ATP12A (the non-gastric P2-type H^+^, K^+^ ATPase), reduction of eotaxin-3 expression due to reduced phosphorylated STAT6 binding, and increased expression of the aryl hydrocarbon receptor (AHR) (9–12). However, the precise mechanisms by which PPI improves pathology in EoE remains unclear.

This study examines the effect of omeprazole, a PPI commonly used for EoE therapy, on the dynamics of esophageal epithelial barrier function and the crosstalk between the epithelium and eosinophils. Using the IL-13-treated air-liquid interface (ALI) culture model, we investigated how omeprazole affects epithelial barrier integrity. We further used an epithelial-eosinophil co-culture system to investigate how omeprazole affects epithelial chemokine secretion and eosinophil activation and migration.

## METHODS

### Esophageal epithelial cell culture

Immortalized human esophageal epithelial cells (EPC2-hTERT, EPC2) were cultured in keratinocyte serum-free media (KSFM, Thermo Fisher Scientific, USA) containing 0.09 mM Ca^2+^, 1ng/ml recombinant human epidermal growth factor, 50 µg/ml bovine pituitary extract (BPE), and 100 U/ml penicillin and streptomycin at 37 °C in a humidified 5% CO_2_ incubator. Cell stimulations were performed in confluent cultures after culturing in KSFM with 1.8 mM calcium concentration for 48 hours to promote maturation and differentiation. Cells were treated with 100 ng/ml recombinant human IL-13 (SRP3274, Sigma–Aldrich) or 50 µM acid-activated omeprazole (O104, Sigma–Aldrich).

### Air liquid interface (ALI) culture

EPC2 cells were grown on 0.4 µm trans wells (Corning Life Science) in KSFM media for three days, then switched to KSFM with 1.8 mM calcium for five days (13). On day 7, media was removed from the upper chamber to promote terminal differentiation. From day 10 to 14, either IL-13 (100 ng/ml), 50 µM acid-activated omeprazole, or a combination of both were added to the basolateral chamber media.

### Bulk RNA sequencing and data analysis

ALI cultures were submerged in TriReagent, and RNA was isolated (Qiagen). RNA sequencing libraries were prepared (Illumina TruSeq RiboZero), followed by paired-end sequencing (Illumina NovaSeq 6000). The raw sequencing data are available in the NCBI GEO series database (14) under accession number GSE273039.

Quality assessment of sequencing reads was performed using FastQC 0.12.0 (1), and contaminant ribosomal RNA was filtered out using FastQ Screen 0.14.1. The subsequent read alignment against GRCh38.p13 reference genome using STAR aligner 2.7.1a. The alignment files were then converted to BAM format, sorted, and indexed using samtools 1.12. Post-alignment quality assessment was executed using Qualimap v2.2.1. Reads were then counted for exonic regions based on gene IDs utilizing the gencode.v29.annotation.gtf annotation file and featureCounts in subread-2.0.2. Summary plots for the output files from the intermediate analysis were generated using MultiQC 1.10. Subsequent downstream RNA-seq analysis was performed using R version 4.3.2.

Genes with counts less than 10 in at least three samples were removed to ensure robustness in the analysis. The gene expression data was normalized for sequencing depth and library composition using variance stabilizing transformation followed by differential gene expression analysis using DESeq2 1.44.0. Principal Component Analysis (PCA) was performed on the 500 most variable genes to visualize sample clustering. The expression patterns were further visualized using a heatmap created with pheatmap 1.0.12. Differential expression analysis involved estimating dispersions and applying Wald tests to determine statistical significance. Volcano plots were used to visualize differential expression across different conditions, and alluvial diagrams depicted gene flow between different conditions, created with ggplot2 3.5.0 and ggalluvial 0.12.5. Pathway enrichment analysis was performed by clusterProfiler 4.12.0, which utilized Fisher’s exact test to identify significant associations with gene ontology biological processes. Statistical significance was determined using the Benjamini-Hochberg (BH) method to correct for multiple testing.

### Transepithelial electric resistance (TEER)

Resistance across the ALI was measured with a Millicell ERS-2 Voltohmeter (Merck Millipore) on days 10, 12, and 14. The TEER for each well was calculated by subtracting the resistance measurement for blank transwells and then multiplying the transwell surface area (0.33 cm^2^ for 24-well plate inserts). Samples with TEER ≤ 200 Ω*cm^2^ on day ten were not used for stimulation experiments.

### Paracellular flux assay

On day 14, 70kDa fluorescein isothiocyanate-dextran (FITC dextran; 3mg/ml; Sigma-Aldrich) was added to the upper chambers and incubated at 37°C for 4 hours. Media from basolateral chambers was transferred to a 96-well plate, and fluorescence was measured at Ex/Em = 485/520 nm (Thermo Scientific Varioskan™ LUX multimode microplate reader). The concentration of samples was calculated using a standard titration curve (30 µg/ml to 0.23 µg/ml).

### ALI histology and immunofluorescence

ALI were harvested on day 14, fixed in formalin, paraffin-embedded, serially sectioned, and stained with hematoxylin and eosin. Images (40X) were analyzed using the Color Deconvolution 2 ImageJ plugin to split the image into separate channels to quantify hematoxylin staining in the basal layer of ALI. A minimum of 5 ALI were analyzed in each condition by examining the mean hematoxylin staining area in a region of interest.

For immunofluorescence, slides were deparaffinized, permeabilized, and blocked in 5% BSA before incubating overnight with mouse anti-DSG1 (1:50, Santa Cruz Biotechnology) at 4°C. Sections were rinsed and incubated for an hour with Donkey anti-Mouse IgG AF488 secondary antibody (Thermo Scientific) and DAPI (Sigma Aldrich).

### Quantitative real-time PCR (qRT-PCR)

RNA was isolated from EPC2 cell cultures, and cDNA was generated from 1 µg of RNA using a high-capacity cDNA reverse transcription kit (Applied Biosystem, CA). 25ng of cDNA was used for qRT-PCR using SYBR fast master mix (Applied Biosystems) and *CAPN14* (BioRad assay ID: qHsaCID0017001) and *GAPDH* (BioRad assay ID: qHsaCED0038674) primers.

### Western Blot

Cell cultures were lysed in RIPA, and 15 µg of protein was loaded into 12% polyacrylamide gels for electrophoresis, then transferred to 0.2 µm PVDF, blocked, and incubated overnight at 4°C in primary antibody (mouse anti-STAT6 1:1000, sc-374021, Santa Cruz Biotechnology; rabbit anti-pSTAT6 1:1000, SAB4300038, Sigma-Aldrich; mouse anti-DSG1, 1:1000, sc-137164, Santa Cruz Biotechnology). Membranes were washed then incubated for 1 hour at room temperature with secondary antibodies (StarBright blue 520 goat anti-mouse IgG and StarBright blue 700 goat anti-rabbit IgG, each at 1:2500, Bio-Rad) and tubulin (hFAB Rhodamine, 1:2500, Bio-Rad). Blot images were acquired using the ChemiDoc Imaging system (BioRad), and analyzed using Image Lab software.

### Eosinophil isolation and culture with EPC2 cells

Whole blood was collected from healthy human subjects who consented to participate. The Children’s Hospital of Philadelphia Institutional Review Board approved the project. Whole blood samples were separated by density gradient, and the pelleted cells were collected. Ammonium chloride solution was used for red cell lysis, and immunomagnetic negative selection was performed to isolate CD45+CD66b+CD16-eosinophils following the manufacturer’s recommendations (EasySep™ human eosinophils isolation kit, STEMCELL). The purity of the isolated eosinophil population was assessed to be >99% of the total CD45+ population using flow cytometry (Supplementary Figure 1).

In eosinophil monocultures, culture media was supplemented with recombinant human IL-5 (10ng/ml, Peprotech) to ensure viability. For coculture experiments, eosinophils (3×10^5^) were added to confluent EPC2 cells in a 24-well plate with a 1:1 mixture of high-calcium KSFM and RPMI (SH30027, Cytiva) supplemented with 10% FBS. Cultures were treated for 24 hours with IL-13, omeprazole, or a combination of both, and then cells and culture supernatants were harvested for downstream analysis.

### ELISA

Cell culture supernatants were collected, and eotaxin-3, CXCL10, periostin, CCL2, and CCL4 concentrations were measured by ELISA (R&D Systems) per the manufacturer’s instructions. The optical density was read at 450 and 570 nm using a multimode microplate reader (ThermoFisher Scientific).

### Eosinophil migration assay

Freshly isolated eosinophils were added to the upper chamber of 3 µm transwell plates (Corning). Conditioned media from EPC2 cultures or fresh culture media supplemented with IL-5 (5 ng/ml) were added to the lower chambers. After 5 hours, eosinophils migrating into the lower transwell chamber were counted.

### Flow cytometry

Eosinophils were collected, washed with PBS, and stained for viability (Zombie NIR™, Biolegend) in PBS for 15 minutes. Cells were washed and incubated with Fc Block (BD Bioscience) for 15 minutes. Cells were stained for BV605 anti-CD45, FITC anti-CD18, BV785 anti-CD11b, and BV421 anti-CD69 (BioLegend). LSR-Fortessa and FlowJo software (BD Biosciences) were used for analysis.

### Statistical analysis

Statistical analyses were performed using a t-test to compare between two groups or a one-way analysis of variance (ANOVA) to analyze multiple groups, followed by post hoc Tukey’s testing to compare the results of multiple groups. *P* values < 0.05 were considered significant. Analyses were performed in GraphPad Prism version 10.2.3.

## RESULTS

### Omeprazole modulates IL-13-induced transcriptional changes in epithelium

To investigate the effect of omeprazole on epithelial transcriptional responses, we used RNA-seq to examine the impact of omeprazole treatment in the IL-13-treated ALI model of EoE (15). Principal component analysis (PCA) of the 500 most variable genes showed that PC1 explained 70% of the variance and differentiated IL-13-treated from non-treated samples. PC2 explained 11% of the variance and distinguished between omeprazole-treated and non-treated samples (Figure 1A, Figure 1B). While IL-13 had the most significant influence on gene expression patterns, omeprazole treatment also affected gene expression.

**Figure 1.**
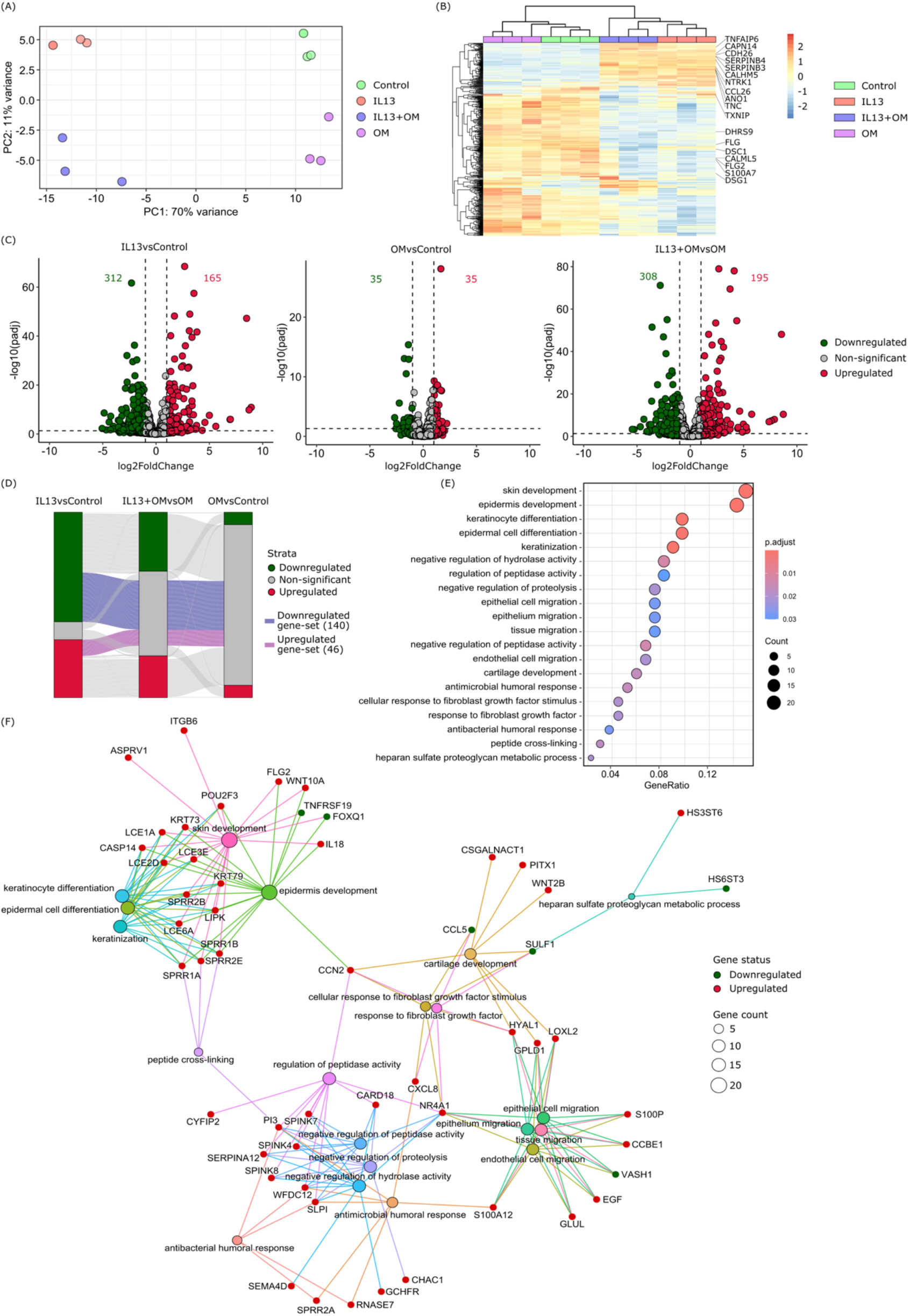
Effect of omeprazole treatment on the differential gene expression of esophageal epithelial cells in IL-13-treated ALI. (A) PCA plot of the top 500 most variable genes. (B) Heatmap of count data for the top 500 most variable genes reveals coherent patterns within each treatment group, (C) Volcano plot illustrating the differential gene expression analysis results as log-fold change versus statistical significance (-log10 adjusted p-value), (D) Alluvial plot represents gene expression changes across treatments. Each stratum in the plot represents treatment conditions, and the widths of the alluvium are proportional to the number of genes in each category. (E) The top 20 most significant gene ontology pathways from analysis of the 140 downregulated and 46 upregulated DEGs in the IL13+OM vs OM comparison (F) Gene-concept network plot of significant genes within the top 20 significant GO-terms. Node size corresponds to gene count, red circles indicate upregulated genes, green circles signify downregulated genes, and edges highlight interactions.

We examined differentially expressed genes using FDR < 0.05 and log2FC ≥ |1| as thresholds. This identified 312 downregulated and 165 upregulated genes in IL-13-treated samples compared to controls (Figure 1C). Consistent with prior studies, IL-13 treatment perturbed genes regulated by STAT6 and associated with EoE pathogenesis, including *CCL26, TNFAIP6, NTRK1, SERPINB4*, and *CAPN14* (Supplemental Table 1)(12). In contrast, omeprazole-treated samples showed fewer differentially expressed genes (35 downregulated and 35 upregulated) than controls, suggesting a less pronounced influence on cellular processes at the transcriptional level. When comparing combined IL-13 and omeprazole treatment with omeprazole alone, 308 genes were downregulated, and 195 genes were upregulated (Figure 1C).

We set out to identify how omeprazole treatment impacts IL-13-regulated genes in esophageal epithelial cells. A subset of genes were downregulated (n=140) or upregulated (n=46) after IL-13 treatment. However, these genes did not show differential expression in ALI cultures co-treated with omeprazole and IL-13 (Figure 1D). This suggested potential gene interactions where omeprazole mitigated transcriptional changes induced by IL-13. We conducted pathway enrichment analysis of these specific genes. Several pathways, including keratinocyte differentiation, epidermal cell differentiation, peptide cross-linking, skin development, and pathways related to inflammation, tissue repair, and cell-cell communication, were significantly enriched in this set of genes (Figure 1E). These findings suggest that omeprazole may modulate pathways associated with inflammation, tissue repair, and cell-cell communication (Figure 1F).

### Omeprazole improves epithelial barrier function in air-liquid interface (ALI) culture

As previously described, IL-13-treated ALI exhibited a significant decrease in TEER on days 12 and 14 (Fig. 2A) and a pronounced increase (∼ 4 fold) in FITC-dextran paracellular flux (Fig. 2B)(16). Histological analysis of IL-13-stimulated ALI demonstrated a reduced differentiated layer and basal hyperplasia (Fig. 2C), indicated by significantly higher basal hematoxylin quantification in the IL-13-treated ALI (Fig. 2D).

**Figure 2.**
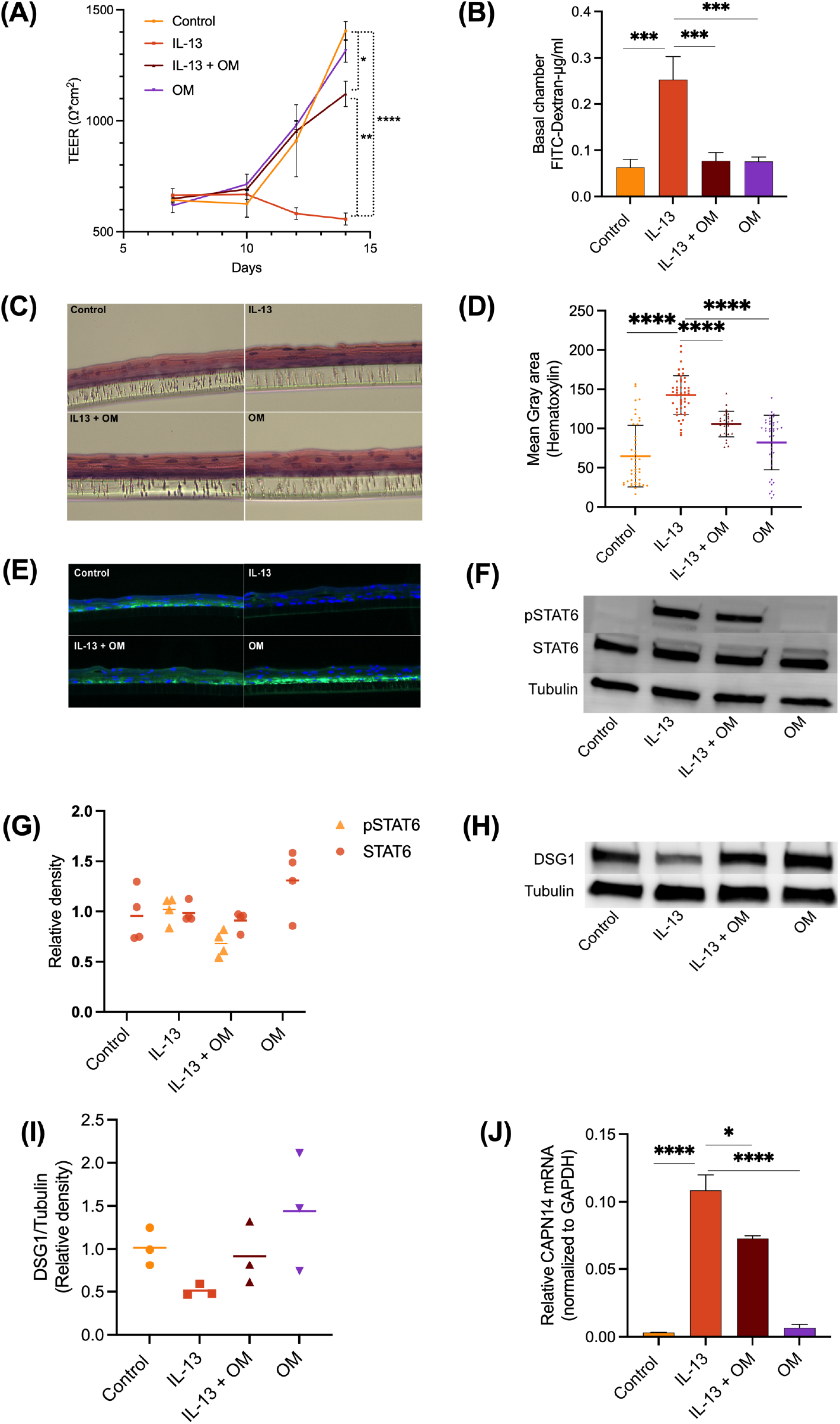
Omeprazole improves esophageal epithelial barrier integrity via regulation of the STAT6 Pathway. (A) Transepithelial electrical resistance (TEER, Ω cm^2^) across the transwell in air-liquid interface (ALI) culture of EPC2-hTERT cells (mean ± SD, n=3 biological replicates, two independent experiments performed). (B) Quantification of translocation of 70kDa FITC-Dextran across ALI membrane into the basolateral chamber (mean ± SD, n=3 biological replicates), (C) Hematoxylin and eosin staining of EPC2-hTERT cells grown in ALI culture, (D) Quantification of hematoxylin staining in ALI (mean ± SD). (E) Immunofluorescence staining of desmoglein-1 (DSG1, green; DAPI, blue) in EPC2 ALI. (F) Representative western blot image of phosphorylated STAT6 (pSTAT6) and total STAT6 in EPC2 (G) Optical density quantification of pSTAT6 and STAT6 relative to corresponding tubulin density (H) Representative western blot image of DSG1 in EPC2 ALI (I) Optical density quantification of DSG1 relative to corresponding tubulin density, (J) Relative mRNA expression of CAPN14 compared to GAPDH (mean ± SD, n=3 biological replicates). *p<0.05, **p<0.01, ***<0.001, ****p<0.0001.

Adding omeprazole treatment to IL-13 stimulation reversed these characteristics of epithelial barrier dysfunction. Although lower than control values, TEER on days 12 and 14 significantly increased compared to ALI treated with IL-13 alone (Fig 2A). There was a similar reduction in FITC-dextran paracellular flux (Fig. 2B), indicating improved barrier integrity. Histological analysis of omeprazole-treated ALI exhibited basal and differentiated morphology similar to control (Fig 2C), and hematoxylin quantification was no different than control (Fig. 2D). These data suggest that omeprazole helps maintain epithelial proliferation and differentiation despite IL-13 stimulation *in vitro*.

### Omeprazole improves barrier integrity by regulating the STAT6 pathway

IL-13 has been shown to upregulate esophageal epithelial calpain-14 (*CAPN14*), leading to loss of desmoglein-1 (*DSG1*) and decreased barrier function (17,18). STAT6 binding in the promoter region of *CAPN14* has previously been shown to regulate calpain 14 expression (19). Therefore, we hypothesized that omeprazole may impact this pathway via effects on STAT6 signaling. To investigate this, we examined the pattern of desmoglein-1 expression in ALI cultures. We found desmoglein-1 staining was decreased in IL-13-treated ALI (Fig. 2E) and further found a 50% reduction in desmoglein-1 protein levels by western blot analysis (Fig. 2H, I). Concurrently, quantification of pSTAT6 and STAT6 demonstrated an IL-13 -mediated upregulation of STAT6 phosphorylation (Fig. 2F, G). Samples treated with both omeprazole and IL-13 exhibited a minimal decrease in pSTAT6 expression compared to IL-13-treated samples, as has previously been described (Fig. 2F, G) (11,20). Desmoglein-1 expression levels in omeprazole-treated groups were similar to those of the control group, regardless of whether IL-13 stimulation was applied (Fig. 2H, I). Similarly, *CAPN14* expression in the IL-13 and omeprazole co-culture group remained upregulated compared to the control but downregulated compared to IL-13-treatment alone (2J).

### Omeprazole downregulates chemokines responsible for eosinophil migration and survival

Prior work has shown that omeprazole impacts pSTAT6 binding at the *CCL26* promoter, downregulating eotaxin-3 in esophageal epithelial cells (11). We examined the overall effect of omeprazole on epithelial chemokine secretion relevant to eosinophil chemotaxis. We performed ELISA to test secreted levels of eotaxin-3, interferon-gamma inducible protein (IP-10/CXCL10), and periostin, associated with eosinophil migration and survival in the epithelium (21–25). IL-13 stimulation upregulated eotaxin-3 expression (106.5 ± 32.62 pg/ml) compared to control (undetected). Eotaxin-3 was not detected in cultures treated with omeprazole alone or the combination of IL-13 and omeprazole (Fig 3A). Similarly, IL-13 stimulation on upregulated CXCL10 expression was approximately 1.5-fold compared to the control. Cultures co-stimulated with IL-13 and omeprazole exhibited slightly decreased CXCL10 expression compared to control and significantly reduced CXCL10 expression compared to cultures treated with IL-13 alone (Fig 3B). Periostin was substantially higher in IL-13 treated (468.9 ± 98.84 pg/ml) compared to control (124.2 ± 25.68 pg/ml), whereas IL-13 and omeprazole cotreated (110.3 ± 14.54 pg/ml) and omeprazole-treated (69.47 ± 23.68 pg/ml) cultures had significantly lower periostin expression compared to IL-13 treated (Fig 3C).

**Figure 3.**
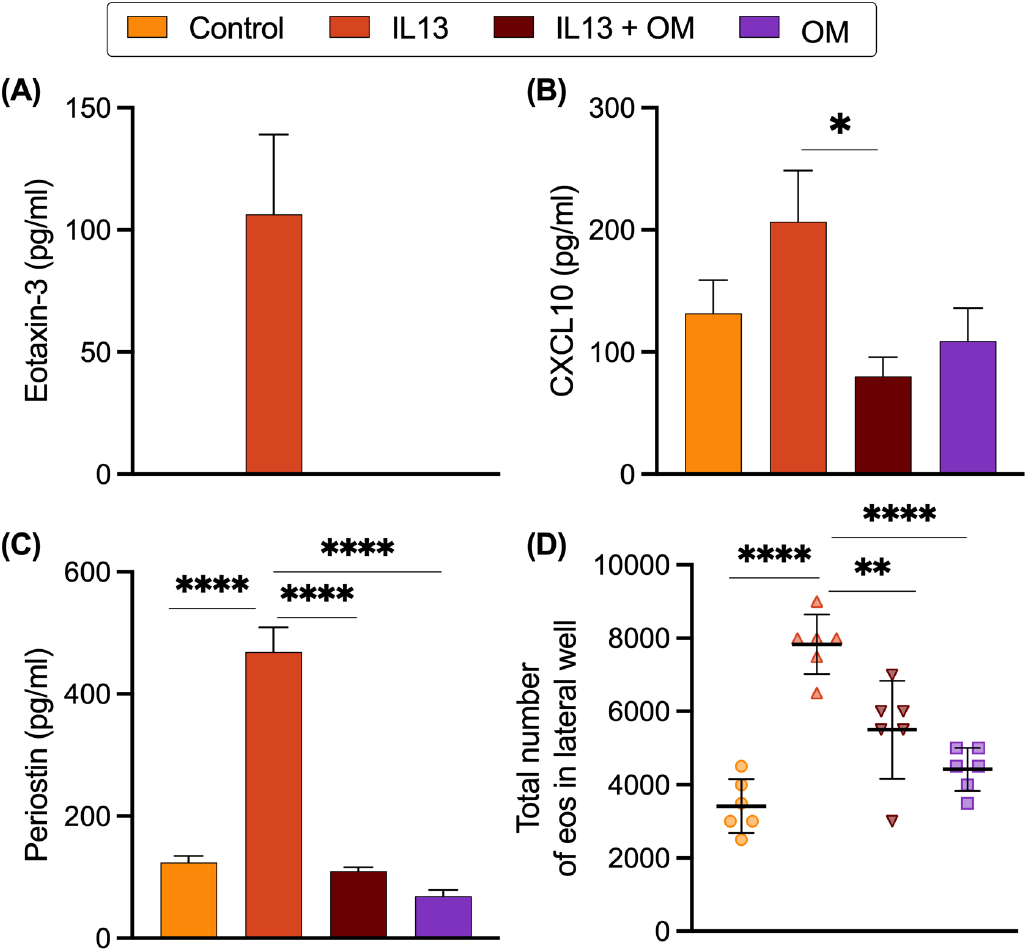
Omeprazole impacts the secretion of inflammatory mediators from esophageal epithelial cells. Levels of (A) eotaxin-3, (B) CXCL10 (interferon gamma-induced protein 10, IP10), and (C) periostin were measured in the epithelial culture media by ELISA. (D) Chemotaxis of eosinophils (eos) through transwells toward epithelial culture-conditioned media in the basolateral chamber. Data are expressed as mean ± SD, n=6, *p<0.05, **p<0.01, ****<0.0001.

Next, we performed an eosinophil migration assay to investigate the net effect of the different chemokine expression profiles. The migration of human eosinophils from normal donor blood was assessed using a transwell assay, and eosinophils migrating through the transwell into the bottom well were quantified. Negligible migration of eosinophils was observed in media containing IL-5, IL-13, omeprazole, or the combination of IL-13 and omeprazole after 5 hours (data not shown). We next assessed eosinophil migration toward conditioned media harvested from EPC2 cultures that had been stimulated in various conditions. Maximum migration of eosinophils to basolateral chambers was observed in response to conditioned media from IL-13-treated cells compared to media from untreated or omeprazole-treated epithelial cells. Eosinophil migration in the omeprazole + IL-13 co-treatment condition was not elevated compared to the control (Fig 3D), consistent with our observations that omeprazole decreases chemokine secretion from epithelial cells.

### Omeprazole downregulates the activation markers of eosinophils

In light of omeprazole’s impact on epithelial chemokine secretion, we hypothesized that omeprazole treatment may impact eosinophil viability and activation due to changes in epithelial mediators. Eosinophils were cultured either with IL-5 alone or in co-culture with EPC2 cells for 24 hours, previously shown to prolong eosinophil survival and enhance viability (26). We assessed eosinophils by flow cytometry to determine viability using a fixable viability dye and including only CD45+ cells (Supplementary Figure 2A). Although IL-5 enhances eosinophil maturation, activation, and survival (27), we observed slightly higher levels of eosinophil survival and activation during co-culture with EPC2 (median 96% vs 67% viability, p=0.029, Supplementary Figure 2B, C). Adding IL-13 or omeprazole to the co-culture did not change eosinophil viability substantially compared to coculture with untreated EPC2 (Figure 4B). We examined eosinophil activation in co-cultures. Eosinophils from co-cultures treated with IL-13 had upregulated CD18 (*p-*value = 0.03), CD11b (*p-*value = 0.061) and CD69 (*p-*value = 0.008) expression compared to eosinophils from nontreated control co-cultures (Fig. 4C). Cocultures treated with a combination of IL-13 and omeprazole did not have upregulation of CD18, CD11b or CD69b (Fig 4C).

**Figure 4.**
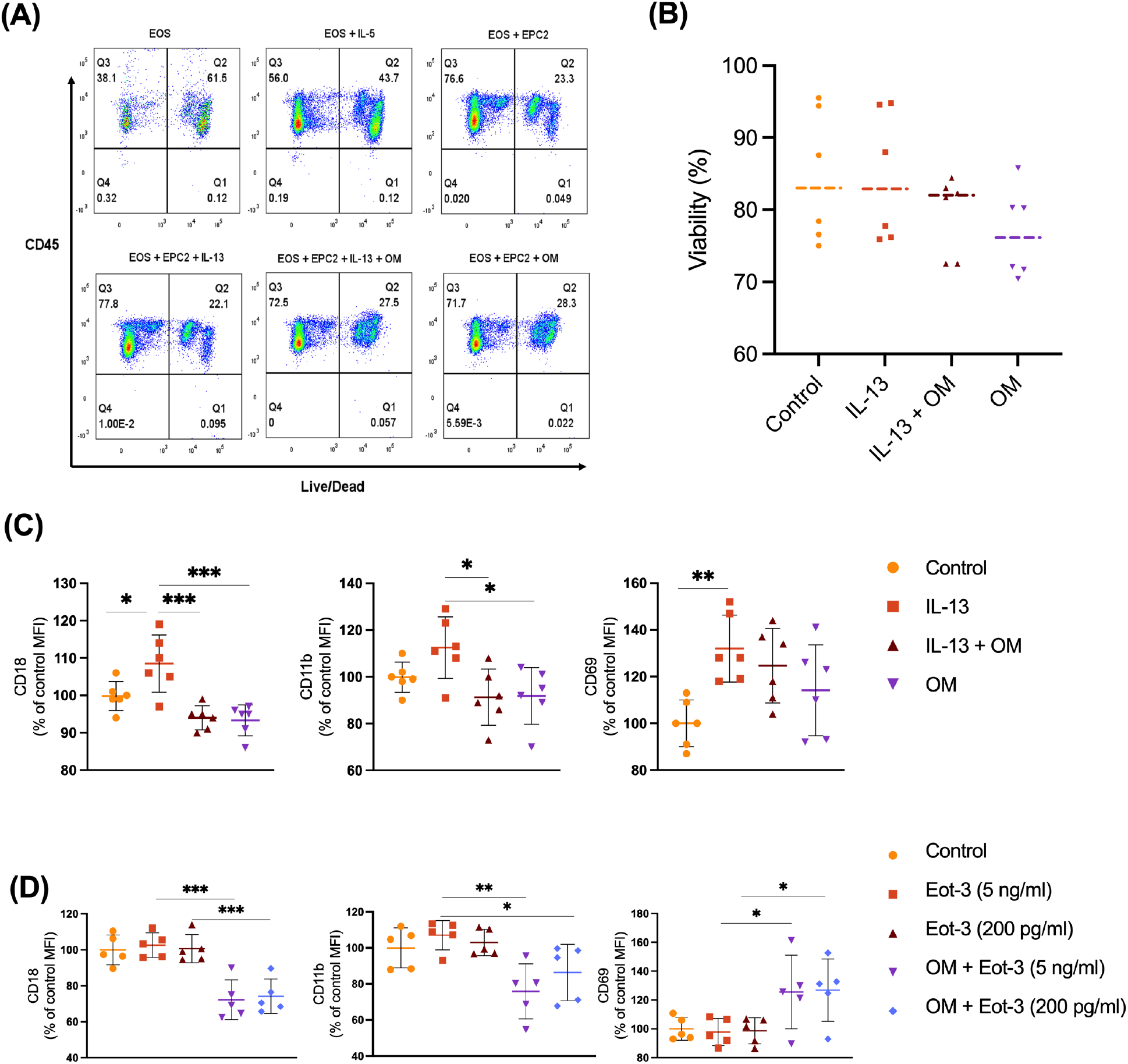
Omeprazole alters the expression of eosinophil activation markers in the eosinophil-epithelial co-culture model. (A) Representative flow cytometry dot plots for eosinophil viability assessment from samples cultured with or without IL-5, or in co-culture with EPC2-hTERT cells (B) Cell viability of eosinophils in co-culture in the presence of IL-13 and/or omeprazole (n=6 biological replicates) (C) Eosinophil activation and adhesion protein expression following omeprazole treatment in the IL-13-treated epithelial-eosinophil co-cultures (mean ± SD, n= 6 biological replicates), (D) Eosinophils were incubated with omeprazole for 30 minutes, then cultured for 24 hours with IL-5 (5 ng/ml) and eotaxin-3 (5 ng/ml and 200 pg/ml) was added as shown to determine if omeprazole affects upregulation of CD11c, CD18 and CD69. Data presented as mean ± SD, n= 5 biological replicates. *p<0.05, **<0.01, ***<0.001.

We next sought to assess if the alterations in eosinophil activation and adhesion molecule expression were due to the direct effects of omeprazole on eosinophils or could be entirely attributed to changing concentrations of chemokines in the conditioned media. We cultured eosinophils without epithelial cells with eotaxin-3 at concentrations of 5 ng/ml or 200 pg/ml to assess the effect of omeprazole pretreatment on activation and adhesion molecule expression. Eosinophils that were pre-treated with omeprazole had decreased CD18 and CD11b and increased expression of CD69 after treatment with eotaxin-3 (Figure 4D).

### Omeprazole attenuates the inflammatory cytokine secretion in Co-culture

To investigate the effect of omeprazole on inflammatory cytokine secretion in a co-culture model, we conducted ELISA assays to quantify soluble inflammatory markers. The overall pattern of eotaxin-3 secretion was similar to that of the epithelial monoculture, as we observed a significant increase in eotaxin-3 production with IL-13 stimulation (54.27 ± 33.83 pg/ml) that was reduced with omeprazole treatment (9.633 ± 11.11 pg/ml) (Figure 5A). Furthermore, IL-13 significantly elevated CXCL10 levels, which was attenuated by co-stimulation with omeprazole or omeprazole alone (Figure 5B).

**Figure 5.**
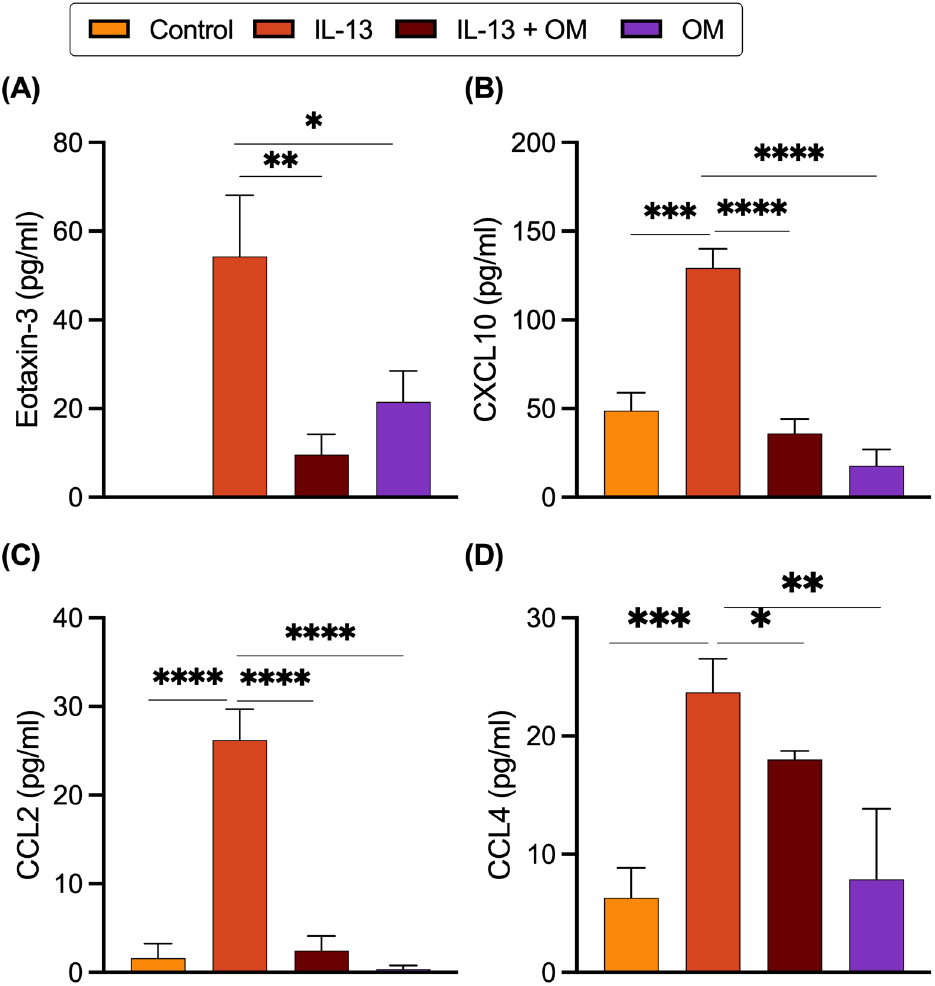
Omeprazole downregulates inflammatory markers levels in IL-13-treated co-culture of eosinophils and EPC2-hTERT. (A) Eotaxin-3 (n=6), (B) CXCL10 (n=5), (C) CCL2 (n=5), (D) CCL4 (n=4). Data are expressed as mean ± SD, and replicates are biological replicates. *p<0.05, **p<0.01, ***p<0.001 and ****p<0.0001.

Although absent in the epithelial cell culture supernatant, quantifiable levels of monocyte chemoattractant protein-1 (MCP-1/CCL2) and macrophage inflammatory protein-1β (MIP-1β/CCL4) were observed in the co-culture, and their expression was significantly upregulated by IL-13. Additionally, treatment with IL-13 and omeprazole, and omeprazole alone led to a significant downregulation of these chemokines in the co-culture system (Figure 5C, D), underscoring the anti-inflammatory potential of omeprazole.

## DISCUSSION

Over recent years, PPIs have become a cornerstone treatment for EoE patients due to multiple factors, including a predictable rate of remission and risk profile, oral bioavailability, and accessibility. Therefore, unraveling the underlying mechanisms driving their efficacy has become increasingly important. In this study of omeprazole in the IL-13 ALI model, our RNA-seq data yielded largely distinct genes regulated by IL-13 and omeprazole. However, approximately 39% of IL-13-responsive genes were modulated by omeprazole treatment. Gene set enrichment revealed that pathways relating to epithelial barrier protein expression and differentiation were enriched among these genes.

Prior work from Zhang et al. suggested minimal change in STAT6 phosphorylation but showed a decrease in STAT6 binding at the eotaxin-3 promoter using ChIP(11). Our data indicated a moderate reduction in STAT6 phosphorylation following omeprazole treatment, and gene expression analysis revealed the potential of omeprazole to regulate several key pathways, including differentiation and barrier protein expression.

Using the ALI model, we demonstrated that omeprazole treatment partially recovered IL-13-induced anomalies in TEER and recovered FITC Dextran permeability and histologic appearance of ALI. This differs from prior observations by Rochman *et al*., where ALI were treated with PPI two days after initiating IL-13 stimulation. In these conditions, no reversal of IL-13-induced barrier dysfunction was observed with delayed onset of PPI treatment. These differences highlight that *in vitro* assays, including the ALI model, are sensitive to conditions such as the timing of stimulation, which is a potential drawback of these models. Clinically, single nucleotide polymorphisms in STAT6 and CYP2C19 have been shown to affect PPI treatment response in patients with EoE (28). These clinical studies highlight the heterogeneity underlying PPI response in EoE patients, which is incompletely modeled using *in vitro* systems like the ALI model.

Our RNA-sequencing data indicated that *CCL26* and *CAPN14* genes were upregulated, and *DSG1* was downregulated in IL-13-treated ALI. Downregulation of *DSG1* has been described in EoE patients in multiple transcriptional studies, and increased expression of the esophageal-specific calpain 14 has been shown to decrease expression of desmoglein-1, contributing to esophageal epithelial barrier dysfunction(16,18). Omeprazole treatment reduced the expression of *CAPN14* and increased the expression of desmoglein-1. Using western blot and immunohistochemistry, we validated the downregulation of desmoglein-1 in IL-13-treated cultures and the normalization of desmoglein-1 expression in ALI following omeprazole treatment. Desmoglein-1 maintains the structural integrity of the epithelium but also inhibits epithelial inflammation via the NF-κB/ERBIN pathway(29).

Here, we show that omeprazole can downregulate the expression and secretion of several key chemokines beyond eotaxin-3. Omeprazole treatment mitigated IL-13-induced expression of eotaxin-3, CXCL10, and periostin, which are pivotal for eosinophil migration and survival within the epithelium(21–25). Multiple studies have shown that epithelial chemokines (i.e., eotaxin-3 and RANTES) or alarmins (i.e., IL-13) upregulate eosinophil expression of CD11b/CD18 (30–32). We demonstrated that the expression of surface markers CD11b, CD18, and CD69 on healthy human eosinophils cocultured with IL-13 stimulated epithelium changed significantly if omeprazole was added. Further, we found a significant decrease in eosinophil migration using conditioned media from EPC2 treated with omeprazole and IL-13 compared to those treated with IL-13 alone.

Recently, several studies have highlighted the role of the esophageal epithelium in providing regulatory signals to eosinophils within the mucosa(26,33). Esophageal epithelial cells have been shown to produce GM-CSF, promoting eosinophil survival *in vitro*. Recent work by Dunn et al. demonstrated that co-culture of eosinophils and epithelial cells in the presence of IL-13 promotes eosinophil survival, upregulates eosinophil activation markers, and promotes a migratory eosinophil phenotype compared to eosinophils from unstimulated co-cultures (33). Our data indicates that omeprazole may impair this process in multiple ways. We observe decreased IL-13-induced chemokine secretion from epithelium in the presence of omeprazole, but there is also a decrease in eosinophil expression of integrins CD18 and CD11b and the activation marker CD69. Together, these data suggest that PPI may work in concert to reduce epithelial secretion of chemokines but also decrease the expression of eosinophil adhesion molecules.

Prior studies in asthma and EoE have shown that the αMβ2 (CD11b/CD18) integrin is upregulated on eosinophils and plays a role in eosinophil migration and tissue persistence during allergic inflammation(34–37). We found both αMβ2 subunits, CD11b and CD18, were upregulated on eosinophils co-cultured with EPC2 compared to those cultured with IL-5 alone. Adding IL-13 to the co-cultures resulted in further upregulation of CD11b and CD18 on eosinophils. Previous studies have shown that αMβ2 binds to periostin, mediating eosinophil chemotaxis into tissue (35). Eosinophils from omeprazole-treated cocultures had decreased expression of these integrins compared to those treated with IL-13 alone. However, neither low (5ng/ml) nor high (200 pg/ml) concentrations of eotaxin-3 increased the expression of CD11b, CD18, or CD69, suggesting that changes in the eotaxin-3 level, as seen in our ELISA data from the culture media, were not entirely responsible for changes in CD11b and CD18 levels. Moreover, omeprazole downregulated CD69 expression in co-culture while increasing its expression in monoculture, suggests an indirect role of omeprazole in downregulating CD69, possibly through the reduction of an epithelial-secreted mediator.

This study demonstrates that omeprazole modulates IL-13-induced transcriptional changes within the epithelium, improves epithelial barrier function, and downregulates key chemokines and activation markers associated with eosinophilic inflammation. These findings provide insight into how PPI therapy may impact crucial pathways and cellular processes implicated in the pathogenesis of EoE. Our data suggest that future research on how PPI treatment modulates epithelial chemokine secretion and eosinophil adhesion molecule expression may further clarify their mechanism in EoE.

## ACKNOWLEDGEMENTS

We gratefully acknowledge the University of Pennsylvania Perelman School of Medicine’s Center for Molecular Studies in Digestive and Liver Diseases (Molecular Pathology and Imaging Core (research Resource Identifier RRID:SCR_022420), and the Children’s Hospital of Philadelphia Flow Cytometry Core and Pathology Core facilities for technical resources. This project was funded via research grants NIH K08AI148456 (to MAR) and the AAAAI Foundation Faculty Development Award (to MAR). The Penn Molecular Pathology and Imaging Core is funded by NIH P30DK0050306. The content in this publication is solely the authors’ responsibility and does not necessarily represent the official views of the National Institutes of Health.

