## Supplementary Figure 1 for "Proton pump inhibitors modulate esophageal epithelial barrier function and crosstalk with eosinophils"

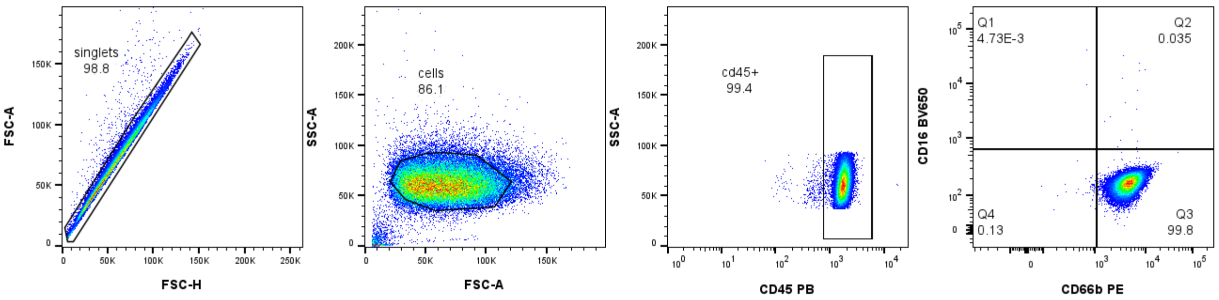


Supplemental Figure 1. The purity of eosinophils (CD16-CD66bCD45+) in the fraction obtained by using the EasySep human eosinophil isolation kit was analyzed by flow cytometry.
