## Supplementary Figure 1 for "Proton pump inhibitors modulate esophageal epithelial barrier function and crosstalk with eosinophils"

(A)

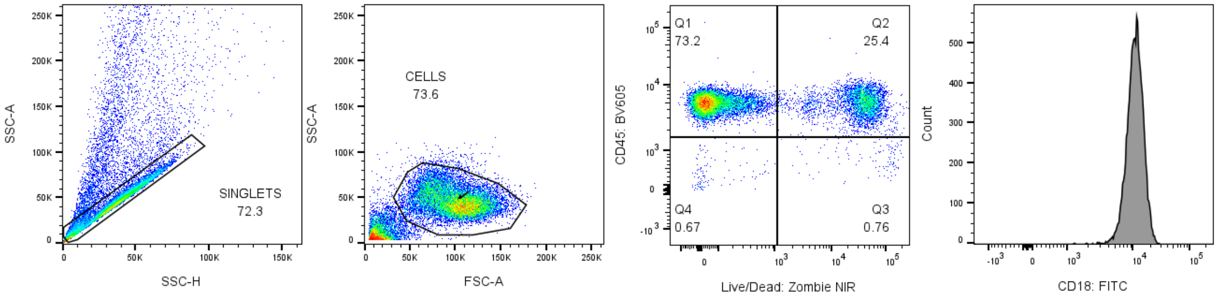


(B)

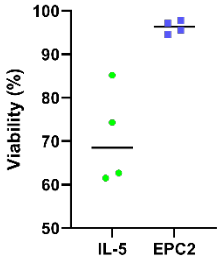


(C)

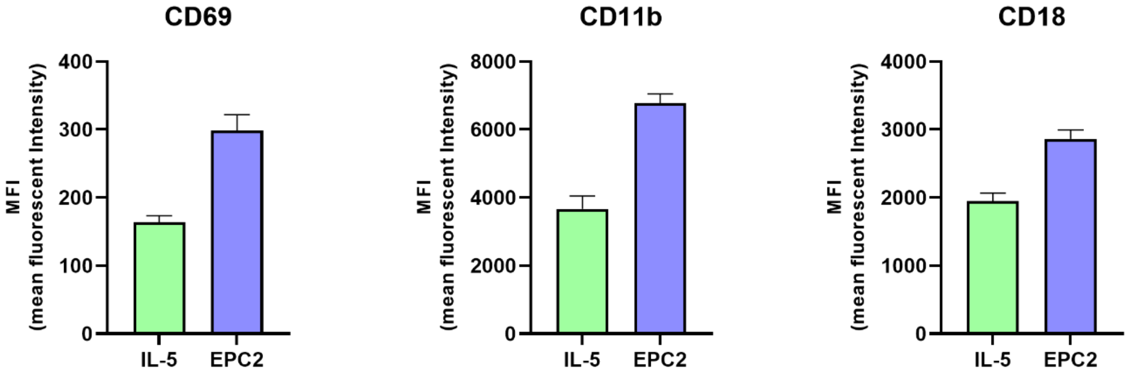


Supplemental Figure 2. Gating strategy and viability of eosinophils in co-culture. (A) Singlets were determined at first and the mean fluorescence intensity (MFI) was determined in live CD45+ cells. (B) percentage of total live eosinophils in culture with IL-5 (10 ng/ml) or EPC2 cells were determined following a similar gating strategy (C) Difference in activation markers of eosinophils in different culture conditions.
